# Relocation of macrophages maintains the barrier function of the urothelium and protects against persistent infection

**DOI:** 10.1101/649137

**Authors:** Jenny Bottek, Camille Soun, Julia K Volke, Akanksha Dixit, Stephanie Thiebes, Anna-Lena Beerlage, Marius Horstmann, Annett Urbanek, Julian Uszkoreit, Martin Eisenacher, Thilo Bracht, Barbara Sitek, Franziska Hoffmann, Nirojah Vijitha, Ferdinand von Eggeling, Daniel R Engel

## Abstract

Macrophages perform essential functions during bacterial infections, such as phagocytosis of pathogens and elimination of neutrophils to reduce spreading of infection, inflammation and tissue damage. The spatial distribution of macrophages is critical to respond to tissue specific adaptations upon infections. Using a novel algorithm for correlative mass spectrometry imaging and state-of-the-art multiplex microscopy, we report here that macrophages within the urinary bladder are positioned in the connective tissue underneath the urothelium. Invading uropathogenic *E.coli* induced an IL-6–dependent CX_3_CL1 expression by urothelial cells, facilitating relocation of macrophages from the connective tissue into the urothelium. These cells phagocytosed UPECs and eliminated neutrophils to maintain barrier function of the urothelium, preventing persistent and recurrent urinary tract infection.

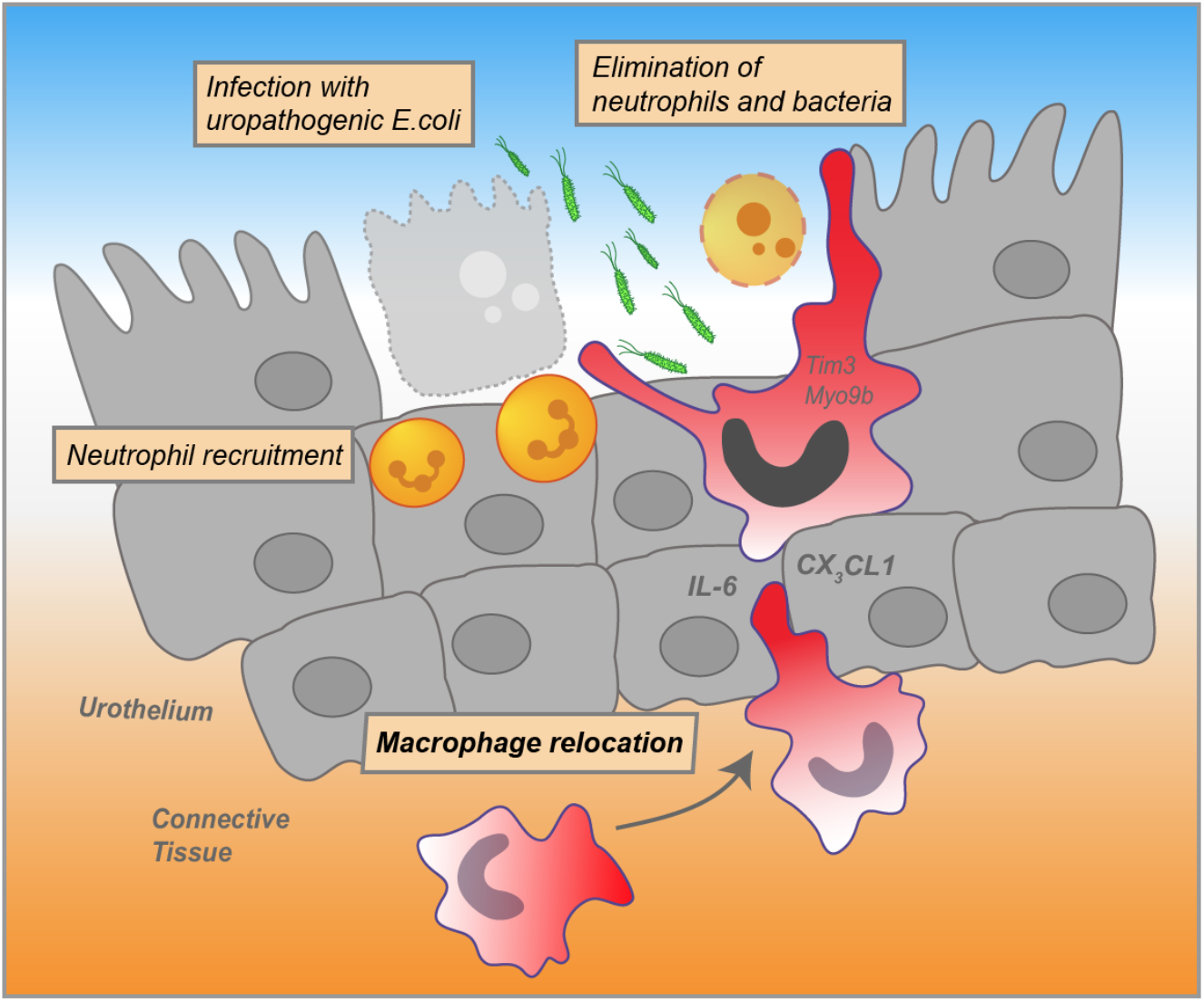
GRAPHICAL ABSTRACT.

## INTRODUCTION

Macrophages are important components of the antimicrobial response and their spatial distribution is crucial to fight off infections (Gordon and Pluddemann, 2017; Regoli et al., 2017). These cells are positioned underneath various epithelia and communication between macrophages and epithelial cells modulates immunity against pathogens (Bhattacharya and Westphalen, 2016; Gross et al., 2015; Wynn and Vannella, 2016). In the urinary bladder, connective tissue macrophages establish a sentinel network, important to maintain homeostasis (Hume et al., 1984; Schiwon et al., 2014). During urinary tract infection (UTI), macrophages contribute to the defense against uropathogenic *Escherichia coli* (UPEC) (Lacerda Mariano and Ingersoll, 2018; Schiwon et al., 2014). UTI is one of the most common and hospital-acquired urothelial infections (Foxman, 2003) with UPECs as the main causative agent (Ulett et al., 2013). The host inflammatory response upon a UPEC infection is critically regulated by macrophages (Abraham and Miao, 2015). These cells sense the pathogen and recruit neutrophils and monocytes to the site of infection by producing chemokines (Schiwon et al., 2014). The absence of macrophages impedes the response against UPECs (Sivick et al., 2010) and there is emerging evidence that macrophages also phagocytose UPECs directly (Mora-Bau et al., 2015).

Imaging approaches, such as state-of-the-art microscopy and mass spectrometry imaging, are powerful technologies to gain spatial molecular information about leukocyte localization in tissues. Matrix-assisted laser desorption / ionization mass spectrometry imaging (MALDI-MSI) is widely used as a label-free imaging technique, which provides important information on the proteomic landscape (Buchberger et al., 2018; Holzlechner et al., 2017), having the potential to unravel molecular mechanisms that regulates the localization of leukocytes (Caprioli et al., 1997; Steurer et al., 2014). Technological and methodological advancements in the past 10–15 years have led to the refinement of this method with a broad range of applications (Oetjen et al., 2016; Seeley and Caprioli, 2008; Urbanek et al., 2016). For proteomic applications (Spengler, 2015), tissue samples are often subjected to an initial on-tissue tryptic digestion step, increasing mass accuracy and resolution. In order to determine the identity of a specific *m/z* value, tandem MS (MS/MS), i.e. LC-MS/MS, in combination with computational methods, existing libraries and coregistration algorithms is able to identify the spatial distribution of proteins (Buchberger et al., 2018; Schey et al., 2013; Vogeser and Parhofer, 2007). Protein Inference Algorithm (PIA) has recently been established to combine peptide spectrum matches from different search engines, increasing consistency of the results (Uszkoreit et al., 2015). This study employs PIA in combination with computational and coregistration methods to establish the coregistration algorithm SPRING (for Spatial PRoteome ImagING), correlating mass spectrometry datasets to extract spatial molecular information in biological samples. SPRING, in combination with state-of-the-art microscopy and experimental *in vivo* targeting approaches unraveled the mechanism and the impact of macrophage relocation into the urothelium during UTI.

## RESULTS

### SPRING identifies macrophage-associated landscapes in the urothelium during UTI

Macrophages establish a dense sentinel network in the connective tissue of the urinary bladder (Schiwon et al., 2014). To study the macrophage response to urothelial infections with uropathogenic *E.coli* (UPEC), we established the algorithm SPRING (Spatial PRoteome ImagING), which employed computational and coregistration methods to analyze mass spectrometry datasets, extracting spatial and molecular information in biological samples. In detail, matrix-assisted laser desorption ionization / mass spectrometry imaging (MALDI-MSI) collected a mass spectrum [m/z] at each pixel on the tissue sections. In order to determine the identity of a specific *m/z* value, liquid chromatography mass spectrometry (LC-MS/MS) was performed and the proteins were detected based on mass matching to databases by the modular pipelining concept KNIME and the Protein Inference Algorithm (PIA) (Uszkoreit et al., 2015). The LC-MS/MS data were matched against a *Mus musculus* FASTA database export together with the common Repository of Adventitious Proteins (cRAP). For each target protein, one shuffled decoy entry was added to estimate the false discovery rate. The spectrum identification was performed using X!Tandem (Craig and Beavis, 2004), the FDR was estimated and filtered on a 1% level, using PIA (Uszkoreit et al., 2015). Thus PIA provided information on the m/z value of the peptides in the LC-MS/MS dataset. For MALDI-MSI, the coordinates and the intensity of all peptides [m/z] for every pixel were extracted after background subtraction and linked to the corresponding peptides and proteins in the LC-MS/MS dataset by SPRING. The expression pattern of all the peptides linked to the same protein were averaged. To study the migration and activation of macrophages in the urinary bladder during UPEC infection, the annotations for the gene ontology (GO) terms “macrophage migration” (GO:1905517) and “macrophage activation” (GO:0042116) were linked to protein accession IDs (Table S1). The expression of the proteins of the GO terms were averaged and a combined distribution map was generated (Figure 1A). SPRING revealed a strong enrichment of the proteins of both GO terms in the connective tissue as well as the urothelium and lumen (Figure 1A), the entry site for invading UPECs. Analysis of the individual proteins of the GO term “macrophage migration” in the urothelium revealed strong expression of Myo9b (Figure 1B), a molecule important for chemokine–induced attraction of macrophages (Hanley et al., 2010). Moreover, the protein Tim3, which contribute to the elimination of apoptotic bodies (Ocaña-Guzman et al., 2016), was strongly expressed in the urothelium (Figure 1B). These data suggest macrophage-specific alterations within the infected urothelium. Indeed, extraction of the 50 highest pixels for the prototypical macrophage marker F4/80 from the urothelial dataset exhibited strong expression of this molecule within the infected urothelium (Figure 1C). Furthermore, Myo9b reached the highest spatial correlation between F4/80 and the proteins of the indicated GO terms (Figure 1D). These data indicate chemokine-mediated relocation of macrophages from the connective tissue into the UPEC–infected urothelium.

**Figure 1.**
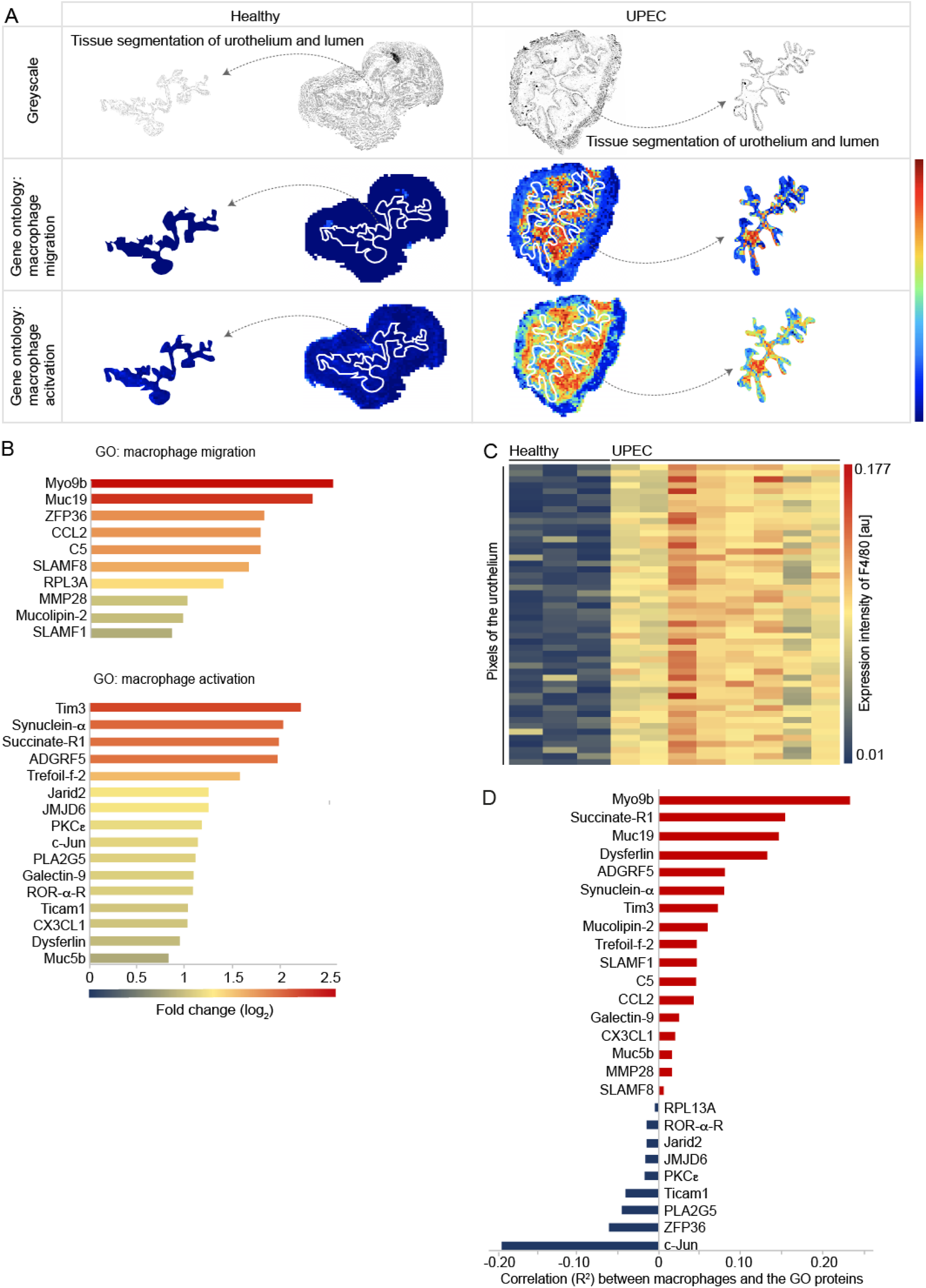
Mass spectrometry imaging indicates macrophage migration and activation within the infected urothelium. (**A**) Representative bladders depicting the spatial distribution of the proteins of the murine GO terms “macrophage migration” (GO: 1905517) and “macrophage activation” (GO: 0042116) in healthy (left) and UPEC–infected (right) conditions by MALDI-MSI. The first row shows a greyscale image and the tissue segmentation of the urothelium from the urinary bladder. The second and the third row shows the expression of the indicated GO terms. The white dashed lines separate the connective tissue from the urothelium and the lumen. The segmented urothelium and lumen are represented as individual images on the right and the left sides. (**B**) Fold changes of the expression intensities of the proteins of the indicated GO terms in the urothelium in healthy versus UPEC-infected samples determined by MALDI-MSI (Formula = Log2(UPEC)-Log2(Healthy)). (**C**) Intensities of the 50 highest-expressing F4/80^+^ urothelial pixels determined by MALDI-MSI in healthy and UPEC-infected bladders. (**D**) Correlation of the expression intensity of F4/80 and the proteins of the GO terms detected by MALDI-MSI. *p < 0.05. Error bars show the mean ± SEM. MP=macrophage; GO=gene ontology; Healthy: n=3; UPEC: n=8.

### Macrophages relocate into the urothelium during UTI

To further study the appearance of macrophages within the urothelium during UTI, tissue sections were analyzed by electron and immunofluorescence microscopy. We found intraurothelial cells with a cellular shape reminiscent of macrophages by electron microscopy (Figure 2A). Immunofluorescence microscopy detected F4/80^+^ macrophages within the EpCAM-1^+^ urothelium (Figure 2B and Figure S2) and these cells were in close proximity to invading UPECs (Figure 2C). To establish an automated and non-biased quantification of macrophages, we developed the algorithm SCHNELL (Statistical Computing on Histology Networks Enabling Leukocyte Location). Macrophage location and abundance were assessed by SCHNELL after segmenting the bladder tissue by the urothelial-specific marker EpCAM-1 and using F4/80 and DAPI for macrophage identification. SCHNELL determined a significant number of macrophages that appeared close to the site of infection (Figure 2D) and an increased density was detected within the urothelium (Figure 2E). Moreover, macrophages were present in the urine already 4 hours after inoculating UPECs into the bladder, corroborating the finding of macrophages relocation during UTI (Figure 2F). These findings confirm the SPRING-based results and demonstrate macrophage accumulation at the site of urothelial infection.

**Figure 2.**
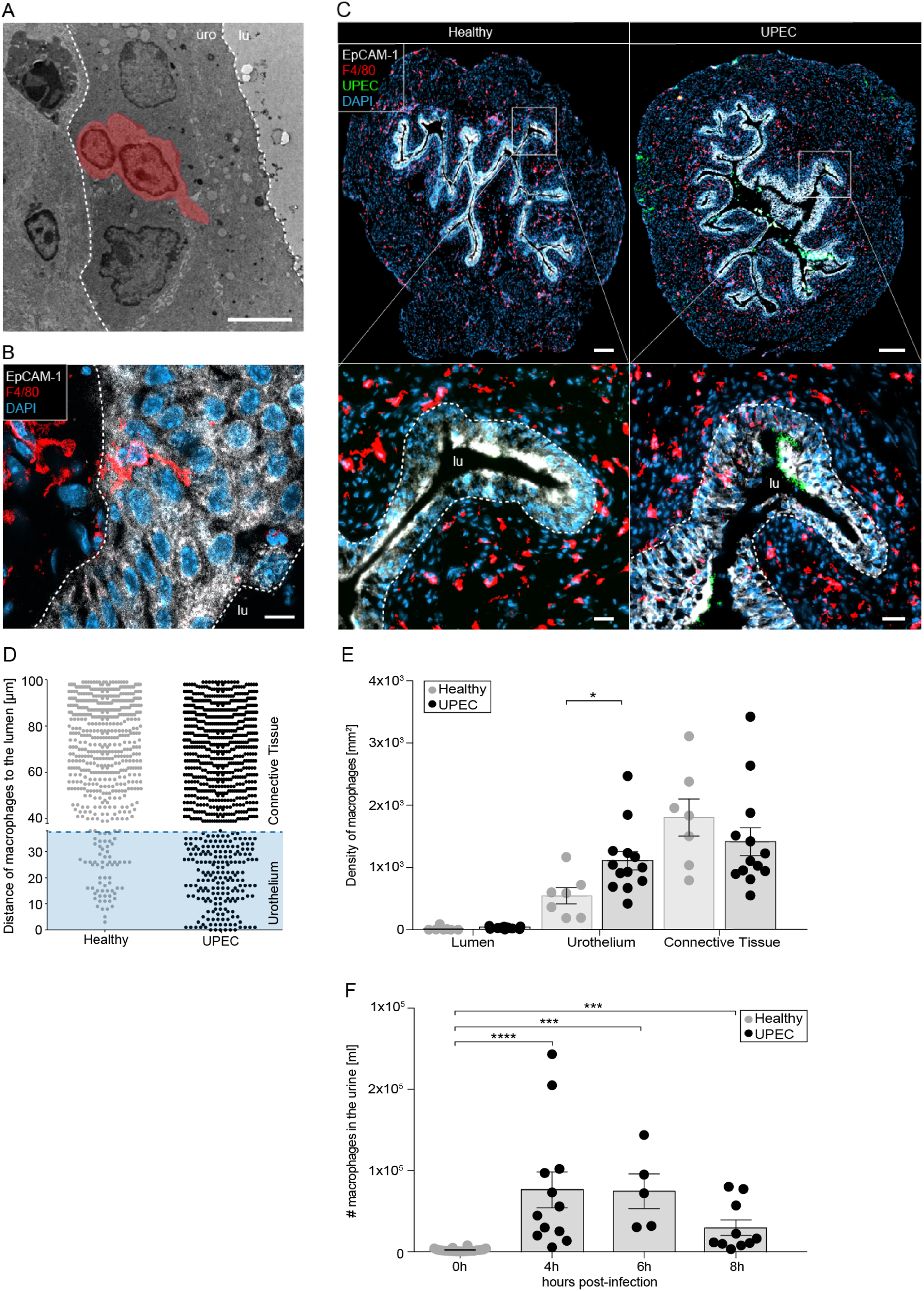
Macrophages accumulate within the infected urothelium. (**A**) Electron microscopy of bladder sections after UPEC infection. The scale bar indicates 10 μm. The macrophage was pseudo-colored based on its cellular structure. (**B**) Detection of intraepithelial macrophages one day after infection by confocal microscopy. The white dashed lines distinguish the urothelium from the connective tissue and lumen. The scale bar indicates 10 μm. (**C**) Immunofluorescence microscopy revealed migration of macrophages towards the infection. The scale bar indicates 200 μm (top images) and 30 μm (bottom images). The white dashed lines represent the border between the urothelium and the connective tissue. (**D**) Representative distance analysis of F4/80^+^ cells. Every dot indicates the distance of a F4/80^+^DAPI^+^ macrophage to the lumen by using the Euclidian Distance map from SCHNELL. The blue box represents the average of the urothelial thickness. Healthy: n=3, UPEC: n=9. (**E**) The density of F4/80^+^ cells was collected by SCHNELL in the various bladder compartments. (**F**) Longitudinal study of F4/80^+^ macrophages in the urine using flow cytometry (0h: n=24, 4h: n=12, 6h: n=5, 8h: n=10). *p < 0.05; ***p < 0.001; ****p < 0.0001. Error bars show the mean ± SEM. Iu=lumen; uro=urothelium.

### Intraurothelial macrophages phagocytose UPECs and eliminate neutrophils

Phagocytosis of pathogens is critical to fight off infections and prevents invasion and persistence of UPECs within the urothelium. We have shown earlier in this study, that Tim3 was strongly upregulated and its expression correlated with the presence of macrophages (Figure 1). Thus, we considered phagocytosis of UPECs as an important defense mechanism, facilitated by relocated macrophages. To study the uptake of bacteria, we inoculated GFP-tagged UPECs into the urinary bladder and imaged intracellular UPECs by confocal microscopy. Macrophages were efficient in phagocytosing UPECs within the urothelium (Figure 3A) and depletion of macrophages by injection of antibodies against the CSF1 receptor (CSF1R) increased the severity of the infection (Figure 3B and Figure S3). Particularly, the infection strength on day 3 was strongly increased after macrophage depletion, indicating persistence and potential hibernation of bacteria within urothelial cells in the absence of macrophages. Using confocal microscopy, we also found Ly6G^+^ neutrophils within intraurothelial macrophages (Figure 3C, 3D and Figure S4) and depletion of macrophages increased the number of neutrophils in the infected bladder (Figure 3E). These data indicate that macrophages are critical to contain the infection and reduce inflammatory molecules from mature neutrophils.

**Figure 3.**
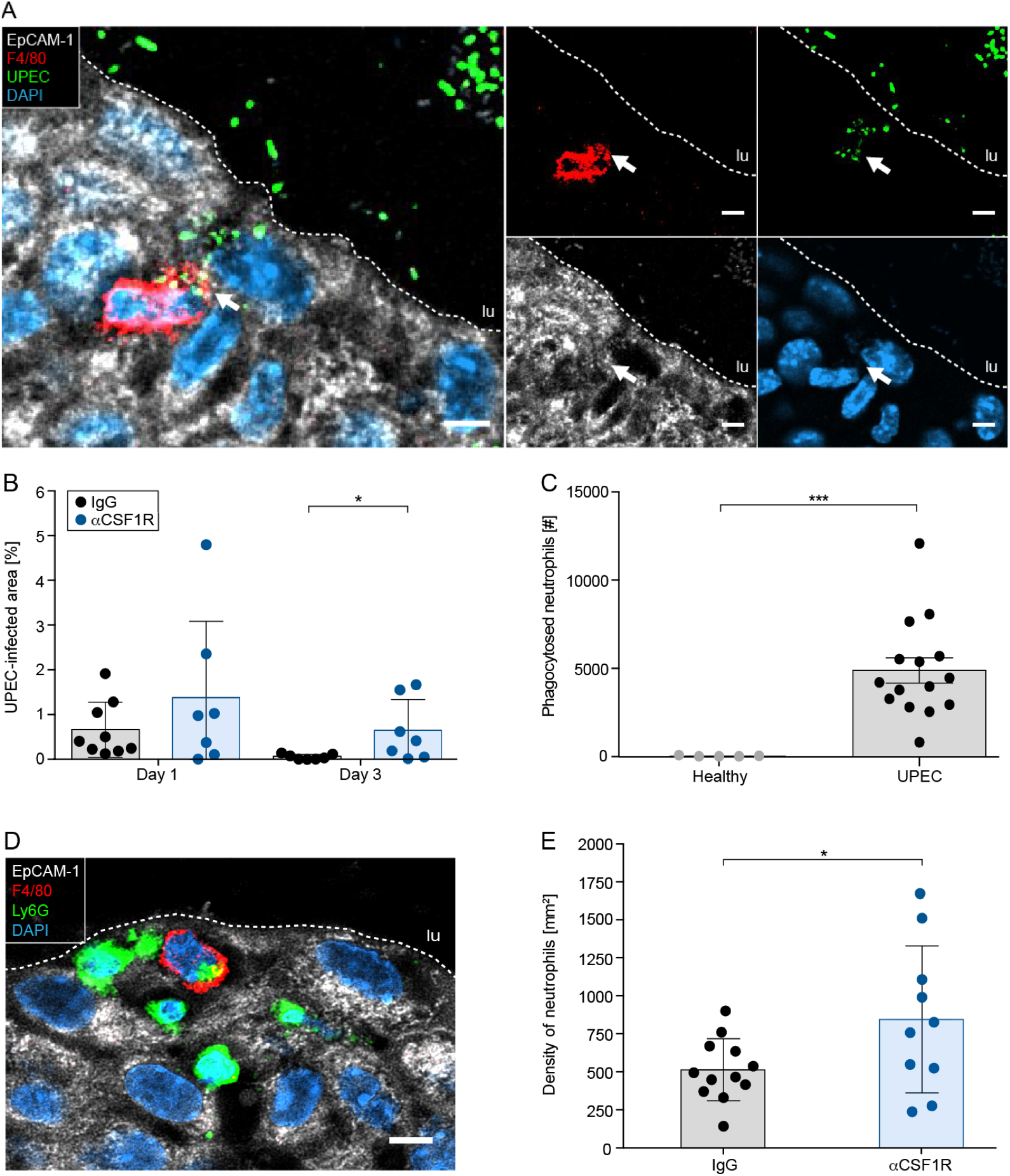
Intraurothelial macrophages phagocytose neutrophils and UPECs. (**A**) Confocal microscopy demonstrating phagocytosis of GFP^+^ UPECs (green) by F4/80^+^ macrophages (red) in the EpCAM-1^+^ urothelium (white). (**B**) Persistence of GFP^+^ UPECs in the urinary bladder determined by fluorescent image analysis (SCHNELL) after macrophage depletion by local inoculation of aCSR1R antibodies. (**C**) Flow cytometry-based quantification of phagocytosis of neutrophils by macrophages (F4/80^+^Ly6G^+^) in bladder digests. (**D**) Confocal microscopy demonstrating phagocytosis of Ly6G^+^ neutrophils (green) by F4/80^+^ macrophages (red) in the EpCAM-1^+^ urothelium (white). (**E**) Density of neutrophils in the urinary bladder after depletion of macrophages by local inoculation of αCSR1R antibodies. *p < 0.05, ***p < 0.001. Error bars show the mean ± SEM. lu=lumen. The scale bars in A and D indicate 5 μm and the white dashed lines distinguish the EpCAM-1^+^ urothelium from the lumen.

### Blockade of IL-6 reduces macrophage migration and aggravates UTI

In order to study the mechanisms that mediate the accumulation of urothelial macrophages, the non-targeted SPRING dataset was analyzed for molecules highly expressed in the urothelium. SPRING revealed strong upregulation of IL-6 (Figure 4A and B) and the expression of this molecule was also strongly detected in the urine post–infection (Figure 4C) and by an antibody-based detection on bladder sections (Figure 4D). To study the relevance of IL-6 for macrophage localization in the urothelium, IL-6 antibodies were topically inoculated into the urinary bladder and the number of macrophages was determined by immunofluorescence microscopy. We found a significant reduction of urothelial macrophages (Figure 5A and B) and an impaired migration to the site of infection (Figure 5C). We also noted aggravated infection levels in the absence of IL-6 (Figure 5D) and impeded expression of the proteins of the indicated GO terms in the urothelium after blocking IL-6 (Figure 5E). These data demonstrate the critical role of IL-6 during UTI and suggest potential proteins involved in macrophage relocation into the infected urothelium.

**Figure 4.**
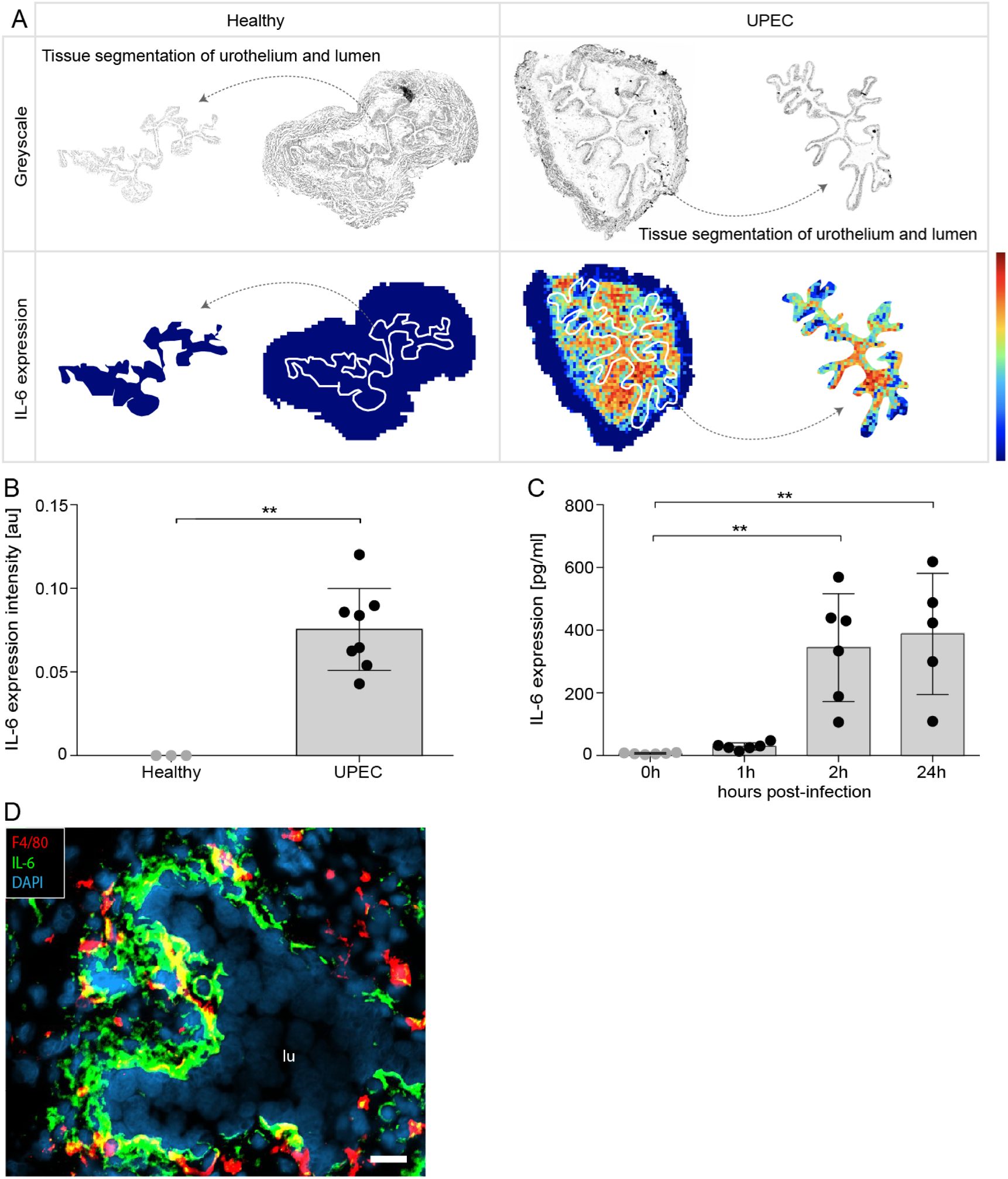
IL-6 is strongly expressed in the urothelium upon UPEC infection. (**A**) Representative spatial distribution of IL-6 in healthy (left) and UPEC-infected (right) bladders by MALDI-MSI. The top row shows a greyscale image and the tissue segmentation of the urothelium from the urinary bladder. The bottom row demonstrates the expression of IL-6. The white dashed lines separate the connective tissue from the urothelium and the lumen. The segmented urothelium and lumen are represented as individual images on the right and left sides. (**B**) Within the urothelium, the number of positive pixels (i.e. whose respective intensity was higher than the background noise) for IL-6 was collected. (**C**) Longitudinal study on the concentration of IL-6 in bladder homogenates (0h: n=6, 1h: n=6, 2h: n=6, 24h: n=5). (**D**) Detection of IL-6 on cryosections 3 hours after infection by immunofluorescence microscopy. **p < 0.01. Error bars show the mean ± SEM. lu=lumen

**Figure 5.**
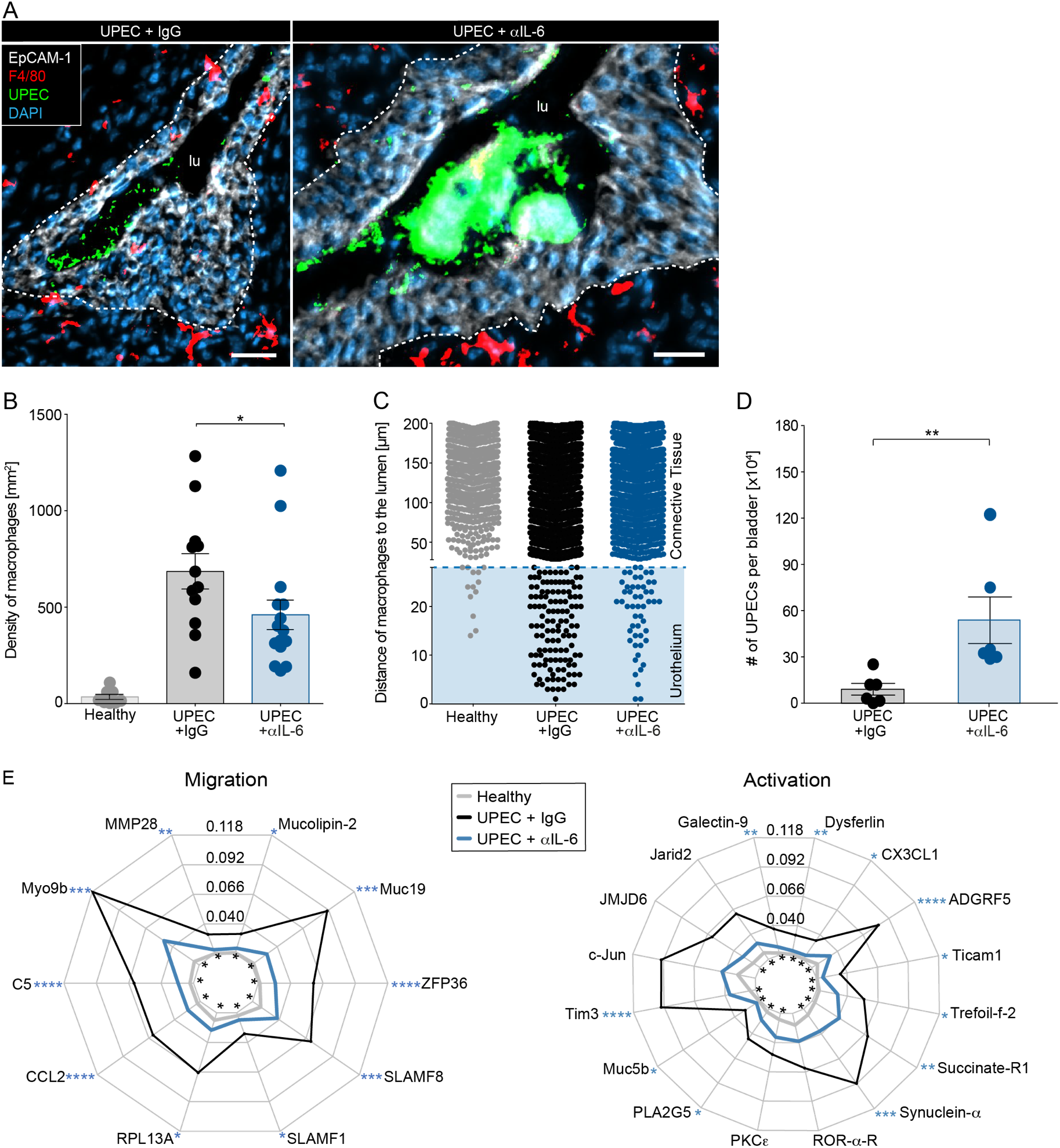
Macrophage relocation and the defense against UPECs depends on IL-6. IL-6 expression was topically inhibited by inoculating IL-6 antibodies (αIL-6). Inoculating IgG served as the isotype control. (**A** to **E**) Mice were infected with GFP-expressing UPECs (green). (**A**) Bladder sections were stained with DAPI (blue), F4/80 (red) and EpCAM-1 (white) and acquired by fluorescence microscopy. Dashed lines indicate the border between the urothelium and the connective tissue. The scale bar indicates 30 μm. (**B**) The density of F4/80^+^ macrophages was calculated by SCHNELL on immunofluorescent images. (**C**) Representative analysis of the distance [μm] of F4/80^+^DAPI^+^ cells to the lumen using a Euclidian Distance map from SCHNELL. The blue box indicates the average of the urothelial thickness. (**D**) The number of UPECs was determined by mechanical disruption of the bladder and culturing the homogenates on *E.coli*-specific agar plates. (**E**) Spider plots of the expression intensities [au] of the GO terms “macrophage migration” (GO: 1905517) and “macrophage activation” (GO: 0042116) in the urothelium determined by MALDI-MSI. Asterisks inside the circle refer to the statistical analysis between healthy and UPEC infected (* in black), whereas the asterisks outside the spider plot (* in blue) compares UPEC versus αIL-6. *p < 0.05, **p < 0.01. Error bars show the mean ± SEM; lu = lumen

### CX_3_CL1 expression depends on IL-6 and mediates macrophage migration into the urothelium

The molecule IL-6 induces classical IL-6 signaling after binding to the IL-6 receptor and recruitment of gp130. Alternatively, IL-6 bound to the cleaved IL-6 receptor activates trans IL-6 signaling via direct binding of the fusion protein to gp130. Macrophages in *LysM^cre/+^ gp130^fl/fl^* animals were still capable to migrate into the urothelium (Figure 6A), indicating that IL-6 acts on urothelial cells to facilitate relocation of macrophages. As presented in Figure 1D, Myo9b showed the strongest correlation with macrophages in the infected urothelium, indicating chemokine-mediated migration of macrophages (Hanley et al., 2010). Among others, CCL1, CCL3, CCL5 and CX_3_CL1 were significantly altered in the absence of IL-6 (Figure 6B). However, inhibition of the corresponding receptors, namely CCR1, CCR3 and CCR5, did not reduce the number of macrophages (Figure 6C). In contrast, targeting CX_3_CR1 impeded macrophage relocation into the urothelium (Figure 6C), demonstrating that IL-6 regulates CX_3_CL1 expression.

**Figure 6.**
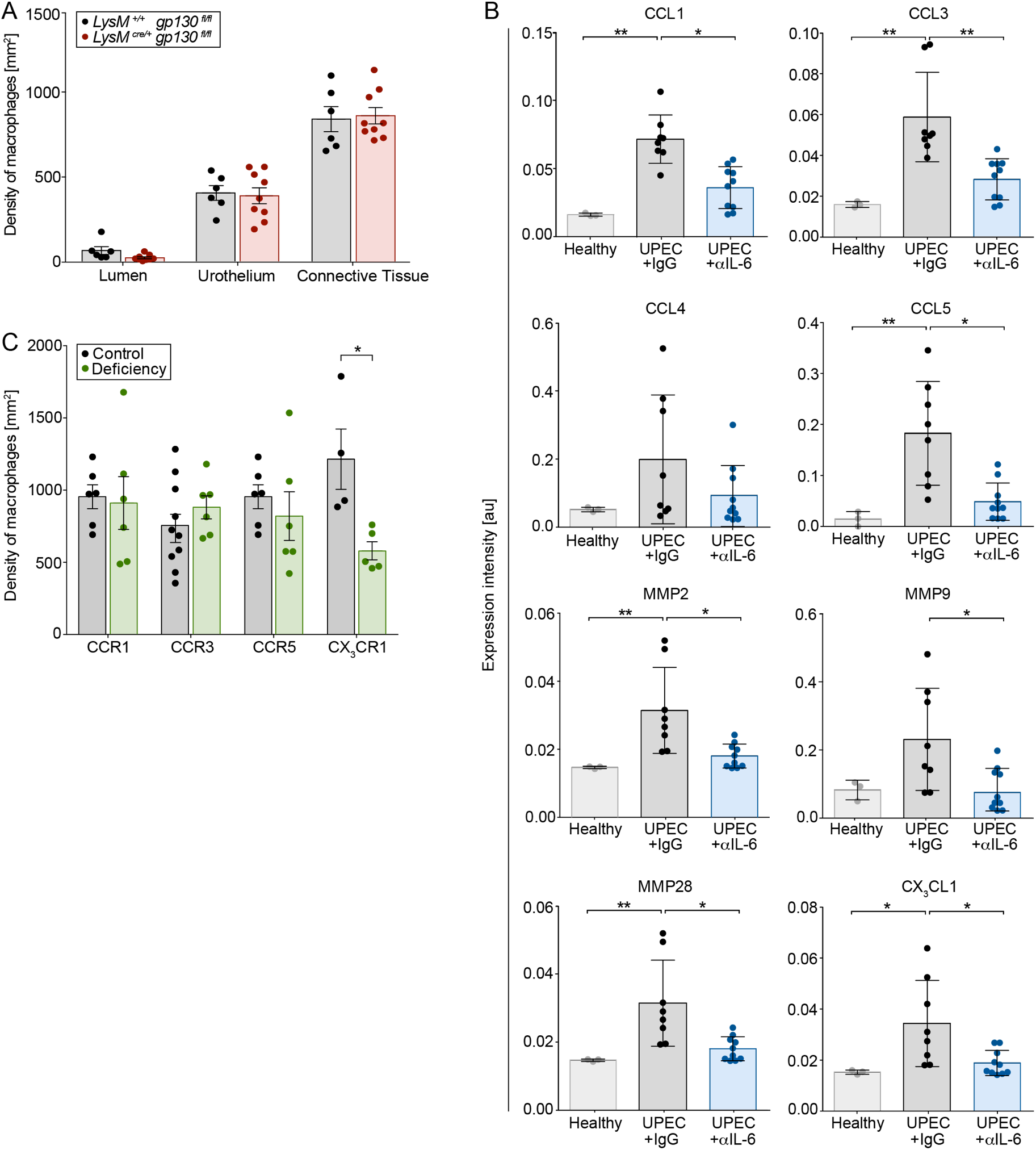
CX_3_CL1 expression depends on IL-6 and mediates macrophage migration into the urothelium. (**A**) The density of F4/80^+^ macrophages, deficient (*LysM^cre/+^ gp130^fl/fl^*) or competent (*LysM^+/+^ gp130^fl/fl^*) in IL-6 receptor signaling, was calculated by SCHNELL on immunofluorescent images. (**B**) Detailed expression intensity [au] of targeted chemokines in the urothelium by MALDI-MSI. (**C**) The urothelial density of F4/80^+^ macrophages was determined on cryosections from fractalkine receptor competent (*Cx_3_cr1^+/gfp^*) and –deficient (*Cx_3_cr1^gfp/gfp^*) mice as well as mice treated with CCR1 (control: DMSO), CCR3 (control: IgG2b) and CCR5 (control: DMSO) inhibitors. *p < 0.05, **p < 0.01. Data are means ± SEM.

## DISCUSSION

Macrophages are found throughout the body, where they have crucial roles in tissue development, homeostasis and remodeling, as well as being sentinels of the innate immune system, contributing to protective immunity (Gordon and Pluddemann, 2017). By employing the novel algorithm SPRING, we now provide evidence that macrophages in the urinary bladder migrate from the connective tissue into the UPEC-infected urothelium in an IL-6–induced CX_3_CL1–dependent manner to contain the infection and eliminate neutrophils, maintaining barrier function of the urothelium and preventing recurrent UTI by hibernating UPECs.

The bladder has an almost impenetrable urothelium, which protects the host tissue from substances that accumulate in the urine and prevents the invasion of microorganisms (Negrete et al., 1996; Sivick and Mobley, 2010). Such physical barrier establishes the first line of defense and responds to potential leaks by producing inflammatory molecules to fight off the infection and recruit leukocytes. A crosstalk of connective tissue macrophages that underlie epithelia has been observed previously (Kopf et al., 2015; Schiwon et al., 2014; Westphalen et al., 2014). The present study demonstrates that signals from the infected urothelium stimulate macrophage relocation from the connective tissue into the infected urothelium. An earlier study detected macrophage protrusions within the urothelium under homeostasis (Hume et al., 1984), indicating that signals derived from the urothelium may induce the recruitment of macrophages into the urothelium. Upon UPEC infection, SPRING colocalized macrophages with Myo9b, an important molecule for chemokine-induced attraction of macrophages (Hanley et al., 2010). These cells phagocytosed UPECs, an important mechanism to contain the infection and prevent hibernation and persistence of UPECs. We speculate that the urothelial barrier function breakdown would occur in the absence of macrophage relocation, leading to recurrent and complicated UTIs. Thus, macrophages within the infected urothelium exert critical anti-microbial functions, potentially through a process recently described as “cloaking” (Uderhardt et al., 2019).

Macrophages in barrier tissues, such as the intestine, lung, skin and urinary bladder, are constantly exposed to environmental cues of the outside world and replenished by invading monocytes during health, inflammation and resolution (Mowat et al., 2017). The differentiation into macrophages is shaped by the tissue microenvironment and migratory and functional adaptations of macrophages are critical to respond to the needs of the anatomical niche. During UTI, large numbers of neutrophils are recruited into the infected urothelium and our data demonstrate that these neutrophils are eliminated by relocated macrophages within the urothelium. Efferocytosis, or elimination of apoptotic cells and bodies, is a physiological and vital mechanism that avoids tissue inflammation and maintains immune tolerance while allowing the development of antibacterial immune responses (Hoffmann et al., 2001). Previously, the transmembrane protein Tim-3 has been shown to capture and eliminate apoptotic bodies after phosphatidylserine union (Ocaña-Guzman et al., 2016). Our data indicate a strong and colocalized expression of Tim3 with urothelial macrophages, suggesting that these cells eliminate mature neutrophils, an important process to reduce inflammatory molecules, preventing inflammation and subsequent tissue damage.

Chemokines are important molecules for macrophage migration. Upon UPEC infection, our algorithm SPRING colocalized macrophages with Myo9b, an important molecule for chemokine-induced attraction of macrophages (Hanley et al., 2010). Indeed, CX_3_CL1 was identified as a crucial chemokine acting as a gate keeper for the entry of macrophages into the infected urothelium. Previously, CX_3_CL1 was critical in intestinal inflammatory disorders and for sampling of bacteria from the intestinal lumen (Niess et al., 2005) and deficiency in CX_3_CR1 increased the translocation of commensal bacteria to the gut draining lymph node (Medina-Contreras et al., 2011). Recently, uptake of UPECs by urinary bladder macrophages has been suggested to impede the adaptive immune response against UPECs (Mora-Bau et al., 2015). Our study demonstrates that CX_3_CR1^+^ macrophages maintain homeostasis and barrier function of the urothelium by phagocytosing UPECs, preventing persistent infection by hibernating UPECs, which subsequently may cause recurrent and complicated urinary tract infection.

The mechanistic investigation of CX_3_CL1^+^–dependent macrophage relocation into the urothelium was facilitated by the coregistration algorithm SPRING (Spatial PRotome ImagING). SPRING was able to generate expression landscapes for proteins of the GO terms “macrophage migration” and “macrophage activation”. Spatial coregistration analysis of the prototypical macrophage marker F4/80 with the proteins of the GO terms indicated a strong increase in the migratory molecules Myo9b and Tim3. Thus, both molecules critically shape macrophage behavior during UTI and our algorithm SPRING was able to identify these changes. Notably, the resolution limit of MALDI-MSI (50μm^2^) might explain the low correlation factors in the linear regression analysis between F4/80 and the proteins of the GO terms. Accordingly, the expression intensity collected in a single pixel might be affected by many different cell types, indicating that technical advancement and increased resolution will be required to achieve cellular resolution. However, segmented analysis of larger specimens, such as the urothelium, solved the cellular resolution limitation and SPRING was able to identify the proteome landscape that facilitated the relocation of macrophages into the urothelium. Importantly, SPRING exploited the label-free MALDI-MSI dataset in a targeted proteomics approach, indicating that the spatial distribution of any molecule and cellular marker of interest could be mapped and colocalized by SPRING in an untargeted manner.

One of the molecules highly expressed in the urothelium upon UPEC infection was IL-6. Pathogen recognition by urothelial cells has been suggested to stimulate IL-6 secretion (Song et al., 2007). Analysis by SPRING revealed strong upregulation of this molecule within the infected urothelium and topical inhibition by local intraurethral administration of antibodies against IL-6 aggravated the infection. Interestingly, the expression of IL-6 strongly correlates with disease severity in humans (Otto et al., 1999) and local production of IL-6 within the urothelium was recently suggested (Dixit et al., 2018). Importantly, application of the JAK-STAT inhibitor Ruxolitinib predisposed patients for UTI (Arana Yi et al., 2015), indicating the critical role of IL-6 signaling for controlling UTI. Analysis of the urothelium environment by SPRING and immunofluorescence microscopy indicated that indeed urothelial cells produced this pleiotropic molecule and topical inhibition of IL-6 within the urinary bladder reduced the chemokines CCL1, CCL3, CCL4, CCL5 and CX_3_CL1. Finally, specific inhibition of the corresponding chemokine receptors identified CX_3_CR1 as most critical for macrophage relocation into the urothelium, indicating that IL-6 shapes the local urothelial microenvironment for CX_3_CL1–dependent macrophage relocation.

In conclusion, our findings support a model in which macrophages maintain barrier function of the urothelium and execute a tissue-protective role in response to bacterial infections. The migratory dynamics of macrophages represent a core function evolved to respond to environmental cues, preventing persistent infection and tissue damage. We conclude that our correlative algorithm SPRING unraveled the mechanism of macrophage relocation during bacterial infections with UPECs. Moreover, such algorithm includes the potential to reveal important cellular and proteomic landscapes in an untargeted approach contained in any dataset. We speculate that intraurothelial macrophages provoke a rather resolving and regenerative environment by reducing inflammatory molecules from mature neutrophils and preventing persistent infections by hibernating UPECs.

## ACKNOWLEDGMENTS

We acknowledge support by the Central Animal Facilities of the Medical Faculty Essen and the Imaging Center Essen (IMCES), in particular Alexandra Brenzel and Dr. Anthony Squire and Dr. Matthias Gunzer for fruitful scientific discussions. This work was supported by grants of the Deutsche Forschungsgemeinschaft to D.R.E (EN984/5-1, EN984/6-1 and SFBTR57), the Marga and Walter-Boll Foundation (220-06-16), Mercur (An-2015-0066) and intramural research funds of the Medical Faculty of the University Duisburg-Essen.

## AUTHOR CONTRIBUTIONS

Investigation, J.B., A.D., S.T., A.L.B., M.H., N.V., J.K.V.; Mass spectrometry measurements, A.U., F.H.; Analysis, visualization and computational analysis, C.S., J.B., B.S., T.Br., J.U., M.E., F.V.E.; Writing, J.B., C.S., D.R.E.; Supervision, D.R.E.; All authors read and commented on the manuscript.

## DECLARATION OF INTERESTS

The authors declare no competing interests.

## MATERIALS AND METHODS

### Animal studies

Female C57BL/6 mice between 6–9 weeks of age were used throughout the experiments. Animals were purchased from Jackson Laboratories or bred and maintained under specific-pathogen-free conditions in the central animal facility at the University Hospital Essen. The following mouse strains were used for the study:

- C57BL/6 J
- F2 crossbreed: *LysM^cre/+^*; B6.129P2-*Lyz2^tm1(cre)Ifo^*/J; Il6st^tm1.1Wme^
- *Cx3cr1^gfp/gfp^*; B6.129P2(Cg)-*Cx3cr1^tm1Litt^*/J

Animal experiments were approved by the local animal review boards (Bezirksregierung Köln, Landesamt für Natur, Umwelt und Verbraucherschutz NRW in Recklinghausen, Germany).

### Urinary tract infection model

Uropathogenic *E. coli* (UPEC) strain 536 (O6:K15:H31) and 536^gfp^ (Engel et al., 2006) were cultured for 3 hours at 37°C in LB medium. Bacteria were harvested via centrifugation at 1500 g for 20 minutes and suspended in 1 ml of PBS. Female mice were anesthetized with a 1:1 mixture of 2% Xylazine and 10% Ketamine. The animals were then infected via transurethral inoculation of 5×10^8^ UPECs in 0.05 ml PBS using a soft polyethylene catheter.

### Blocking experiments and macrophage depletion

Blocking experiments were performed by transurethral injection of 1.5 μg/g mouse weight of the indicated antibodies and inhibitors (key resource table) into the bladder lumen 1 hour post–infection. Anti-CSF1R (αCSF1R) antibody was used to deplete bladder macrophages. Animals received two intraperitoneal injections of αCSF1R antibody or isotype control (20 μg/g mouse weight on day 1 and 10 μg/g mouse weight on day 2).

### Isolation of leukocytes from the urinary bladder

Bladders were sliced into small pieces using a scalpel and then digested for 45 minutes at 37°C with 0.5 mg/ml collagenase and 100 mg/ml DNAse I in RPMI 1640 medium supplemented with 10% heat–inactivated FCS, 20 mM HEPES, 1 mM L-glutamine and antibiotics. Single-cell suspensions were filtered through a 100 mm nylon mesh and analyzed by flow cytometry.

### Flow cytometry and microscopy

#### Cell surface staining

Single-cell suspensions were washed with PBS containing 0.1% BSA and 0.1% NaN_3_, and Fc-receptors were blocked with human immune globulin. Titrated amounts of the fluorochrome–labeled antibodies were used for staining. Cells were measured using an LSR Fortessa (BD Biosciences) and analyzed with the FlowJo software 10. Absolute cell numbers were calculated by adding a fixed number of APC–labeled microbeads (BD Biosciences) to each sample.

#### Electron microscopy

Bladder tissues were fixed with 3% glutaraldehyde in 0.1 M cacodylate buffer [pH 7.4] followed by 2% osmium tetroxide. Tissues were embedded in Epon 812 embedding resin, and 40 to 50 nm thin sections were cut with an LKB ultramicrotome UM IV (Leica) and analyzed using a CM10 electron microscope (Philips).

#### Immunofluorescence microscopy

Bladders were fixed overnight in PLP buffer [pH 7.4, 0.05 M phosphate buffer containing 0.1 M L-lysine, 2 mg/ml sodium periodate, and paraformaldehyde with a final w/v concentration of 4%]. Subsequently, bladder tissues were equilibrated in 30% sucrose for 24 hours and then frozen in Tissue-Tek OCT. Bladder sectioning was performed at −20°C by using a cryostat. Sections with a thickness of 10 μm were rehydrated using PBS containing 0.05% Triton X-100, blocked for 1 hour with PBS containing 1% BSA and 0.05% Triton X-100, and 2-3 sections per bladder were imaged on a Zeiss AxioObserver.Z1 or Leica SP8 gSTED Super-Resolution confocal and FLIM. For IL-6 immunostaining, animals were sacrificed 3 hours post–infection and isolated bladders were pre–incubated with 1 μg/ml Golgi-Plug in RPMI [10% FCS, 1% L-glutamine, 1% Penicillin/Streptomycin) for 4 hours at 37°C, snap-frozen in Tissue-Tek OCT and processed for cutting as described above.

### Key resource table

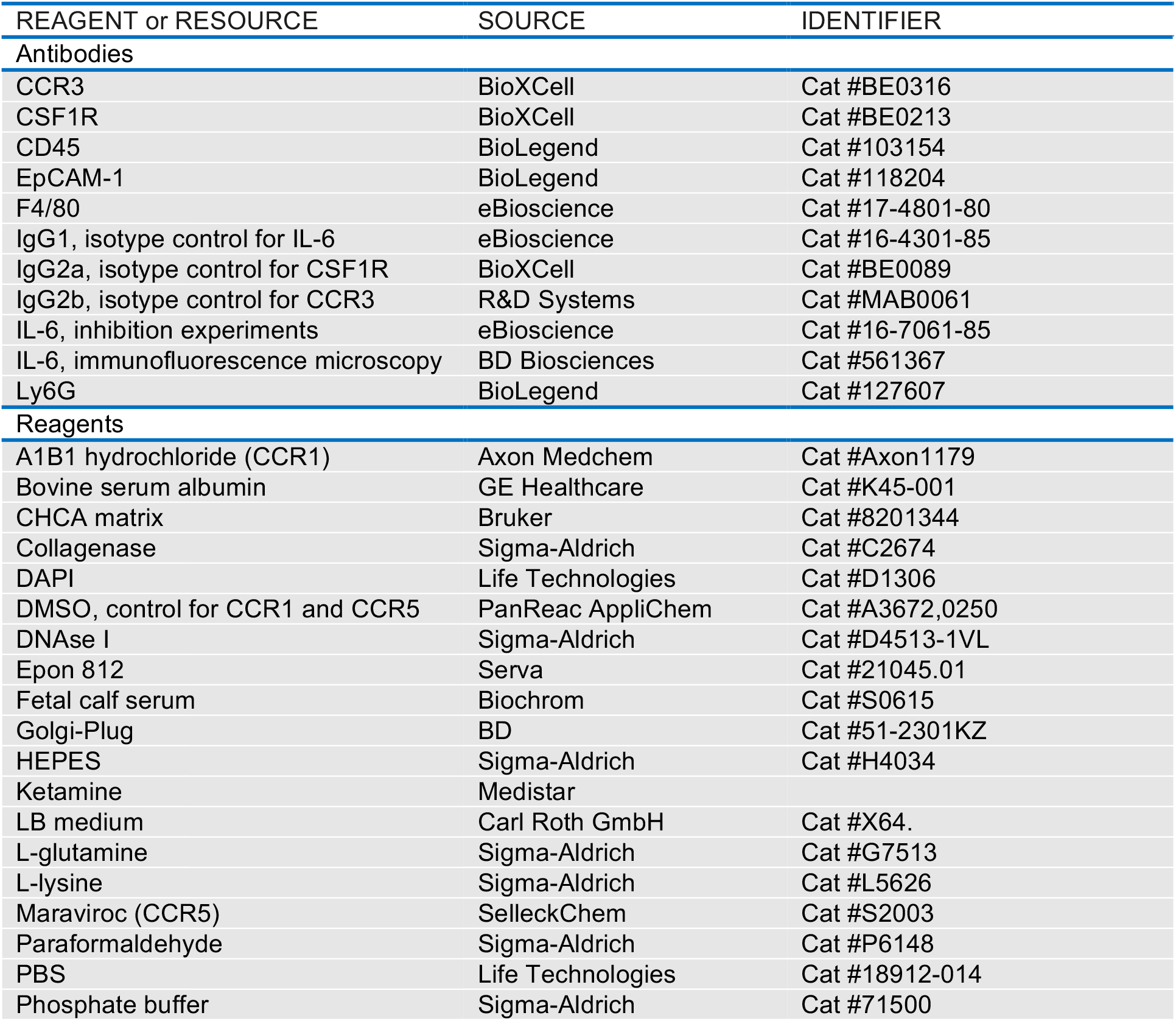
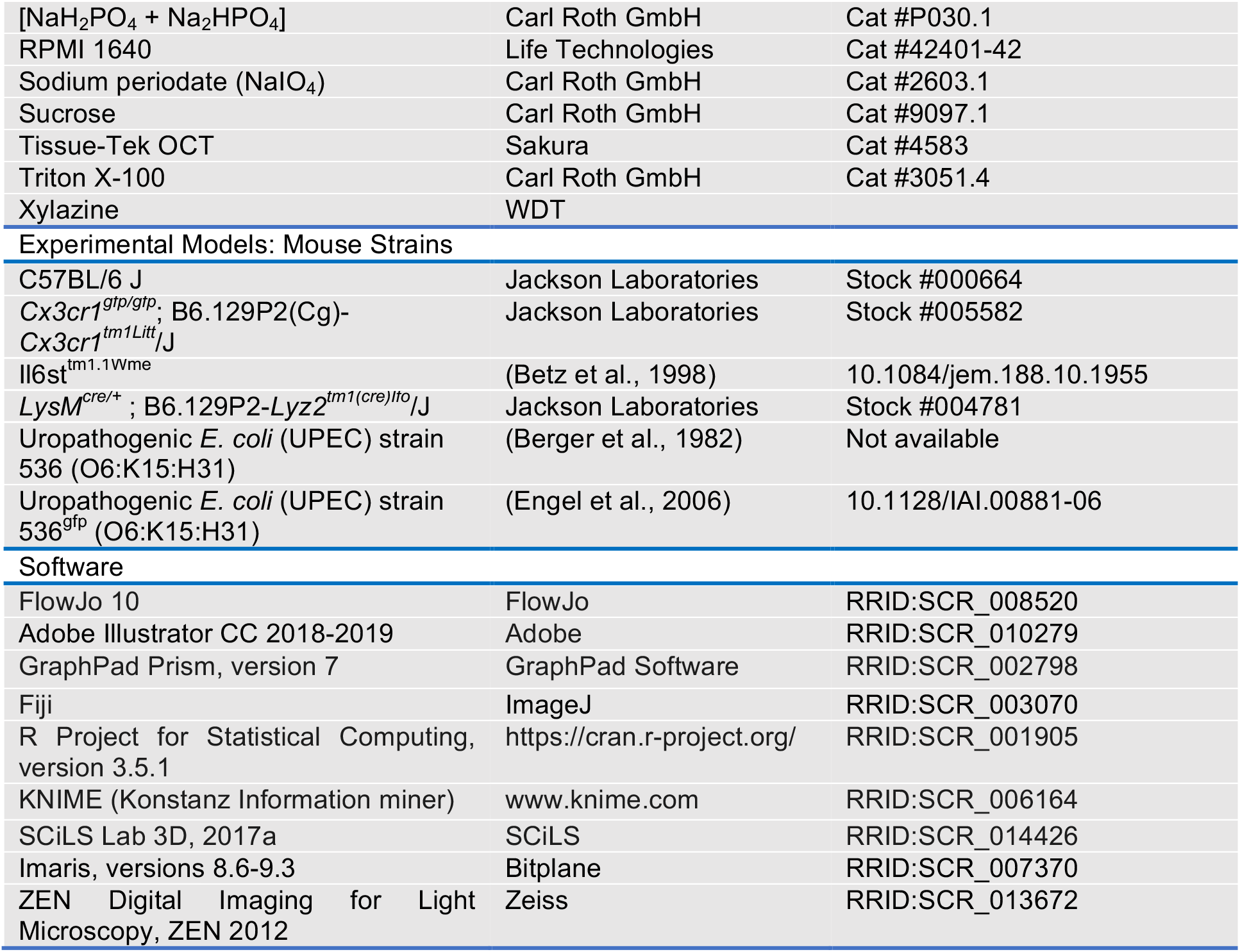

### Analysis of microscopy images by SCHNELL (Statistical Computing of Histology Networks Enabling Leukocyte Location)

An intensity threshold was used to generate masks for each fluorescent channel and the binary information for cellular and nuclear signals was coregistered. Automated analysis of cell densities was performed by a Java based algorithm SCHNELL (Statistical Computing of Histology Networks Enabling Leukocyte Location). By using ImageJ, overlapping mask regions were used to identify cells, which were marked with a point at the center of the DAPI^+^ cell nucleus. The bladder tissue was segmented into lumen, urothelium and connective tissue by employing the EpCAM-1 signal and cell densities were calculated. The distance of macrophages to the infection was assessed, using the lumen mask as the target. The Euclidian Distance was calculated on 16-bit images by shading the pixels from luminal to abluminal. The Euclidian Distance map was overlaid with the macrophage point map in order to calculate the distance of macrophages. The distances were converted from pixels to microns in R using the following formula:

Distance[μm] = Distance[pixels] x (Size of 1 charged-coupled device element[μm]/magnification)

### Matrix-assisted laser desorption / ionization imaging (MALDI-MSI)

#### Sample preparation

Bladders were perfused transcardially with a total volume of 20 ml PLP buffer [pH 7.4, 0.05 M phosphate buffer containing 0.1 M L-lysine, 2 mg/ml sodium periodate, and paraformaldehyde with a final w/v concentration of 4%] at a perfusion rate of 4 ml per minute. Perfused bladders were fixed in PLP buffer overnight. Fixed bladders were equilibrated in 30% sucrose for 24 hours and 5 μm sections were cut at −20°C using a cryostat. The slices were mounted on indium tin oxide (ito) slides. Tissue sections were washed with ethanol (70% and 95%) for 2 minutes. The following steps of on-section tryptic digestion and application of CHCA matrix were performed as described previously (Hoffmann et al., 2019).

#### Data acquisition and analysis

Measurements were performed using the UltrafleXtreme mass spectrometer from Bruker. The parameters were as follows: reflective positive ion mode, mass range: 700-4000 m/z, spatial resolution: 50 μm^2^, laser beam size: medium, shots per position: 300-500, sampling rate: 2.5 GS/s. For external instrument calibration the Bruker peptide calibration standard was spotted next to the tissue section. The SCiLS Lab software was used for data and imaging analysis of the MSI measurements. All data were total ion counts (TIC), normalized and baseline removed.

### Liquid chromatography mass spectrometry (LC-MS/MS)

#### Sample Preparation

Tissue sections were scraped off the glass slides and lysed by addition of 20 μl ammonium bicarbonate (50 mM), containing 0.1% RapiGestSF surfactant per sample. Disulfide bonds were reduced with 5 mM dithiothreitol at 60°C for 30 minutes and alkylated with 15 mM iodoacetamide for 30 minutes at ambient temperature in the dark. Lysed proteins were digested with trypsin overnight at 37°C using 0.1 μg trypsin per sample. For acidification, trifluoroacetic acid (TFA) was added (0.5%, 30 minutes, 37°C) and the samples were centrifuged (10 minutes, 16,000 g) for removal of precipitated RapiGest. The supernatants were collected, dried in a vacuum centrifuge, and dissolved in 50 mM ammonium bicarbonate. For peptide purification, Sera-Mag beads were equilibrated with water and peptides were added to the beads (40 μg beads per sample, hydrophilic and hydrophobic beads mixed 1:1), before acetonitrile (ACN) was added to a final fraction of 95%. Peptides were allowed to bind for 10 minutes at room temperature before the beads were immobilized using a magnetic rack. Beads were washed with ACN two times, dried and 20 μl 0.1% TFA per sample were added. The samples were sonicated on ice for 5 minutes and the eluted peptides were separated from the beads using a magnetic rack.

#### LC-MS/MS analysis

The LC-MS/MS analysis was performed using an Ultimate 3000 RSLCnano system coupled online to an Orbitrap Elite mass spectrometer (both Thermo Scientific). Peptides dissolved in 0.1% TFA were pre–concentrated on a C18 trap column (Acclaim PepMap 100; 100 μm × 2 cm, 5 μm, 100 Å; Thermo Fisher Scientific) within 7 minutes at a flow rate of 30 μl/min with 0.1% TFA. Subsequently, the peptides were separated on an analytical column (in-house packed C18 analytical column, ReproSil®-Pur (Dr. Maisch HPLC GmbH, Ammerbuch, Germany), 75 μm × 40 cm, 1.9 μm, 120 Å) by a gradient from 5% to 40% solvent B over 98 minutes (solvent A: 0.1% FA, solvent B: 0.1% FA, 84% ACN; flow rate 300 nl/min; column oven temperature 65°C). The instrument was operated in a data–dependent mode. Full-scan mass spectra in the Orbitrap analyzer were acquired in profile mode at a resolution of 60,000 at 400 m/z and within a mass range of 350–2000 m/z. MS/MS spectra were acquired in data–dependent mode at a resolution of 5,400. For MS/MS measurements, the 20 most abundant peptide ions were fragmented by collision–induced dissociation (CID, NCE of 35) and measured for tandem mass spectra in the linear ion trap.

### Spatial PRoteome ImagING (SPRING): An algorithm combining computational and coregistration methods to analyze mass spectrometry datasets

SPRING combines LC-MS/MS and MALDI-MSI datasets to generate tissue proteome landscapes. The LC-MS/MS data were matched against a *Mus musculus* FASTA database export of UniProt (UniProt, 2019) (release 2018_06, 73045 entries) together with the common Repository of Adventitious Proteins (cRAP, 115 entries). For each target protein, one shuffled decoy entry was added to estimate the false discovery rate. The spectrum identification was performed using X!Tandem (Craig and Beavis, 2004), the FDR was estimated and filtered on a 1% level using PIA (Uszkoreit et al., 2015). PIA provided information on the m/z value of the peptides, including the corresponding protein. For MALDI-MSI, the coordinates and expression intensities of all peptides [m/z] per pixel were extracted by SCiLS Lab after background subtraction. These values were correlated to the corresponding peptides [m/z] in the LC-MS/MS dataset. For proteins linked to more than one peptide, the distribution per pixel was averaged in order to extract one spatial distribution map per protein and sample and spatial distribution maps were generated.

### Statistical tests

Results are presented as means ± SEM and p-values are depicted in the figures. Unpaired Mann-Whitney (two-tailed) or Kruskal-Wallis tests were performed to compare two or more groups. Where applicable, multiple comparison post-hoc corrections were applied (Dunn’s for multiple groups). Paired data from the longitudinal experiments were analyzed using Friedman’s rank test.

## SUPPLEMENTAL INFORMATION

**Table S1.**
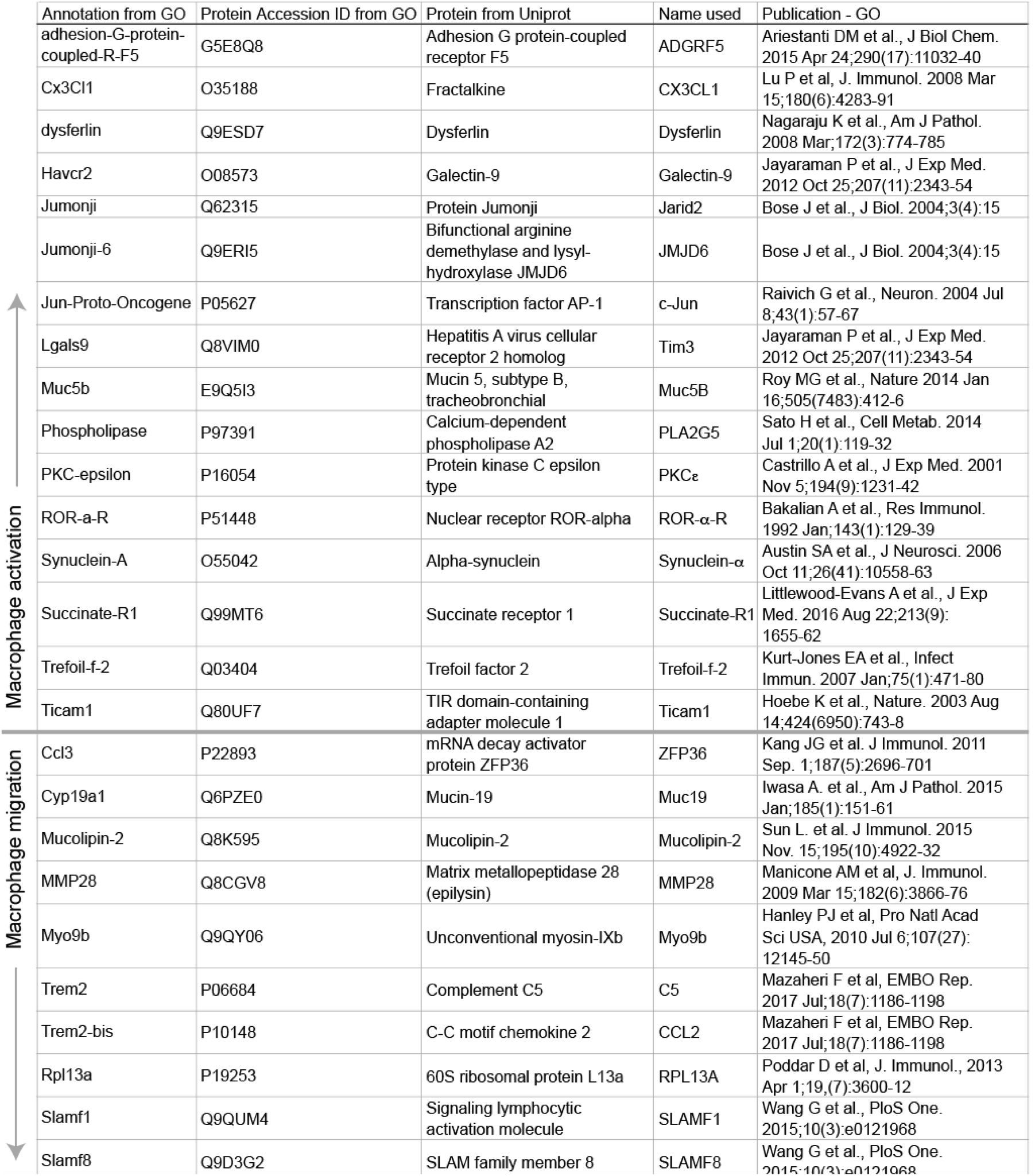
Translation of the murine genes of the gene ontology terms (GO) “macrophage migration” (GO:1905517) and “macrophage activation” (GO: 0042116) into the corresponding proteins. Related to Figure 1 and 5. Annotations of the indicated GO terms were linked to protein accession IDs, which were used to identify the corresponding protein in the protein-database Uniprot. The protein names given by Uniprot were used throughout the manuscript.

**Figure S2.**
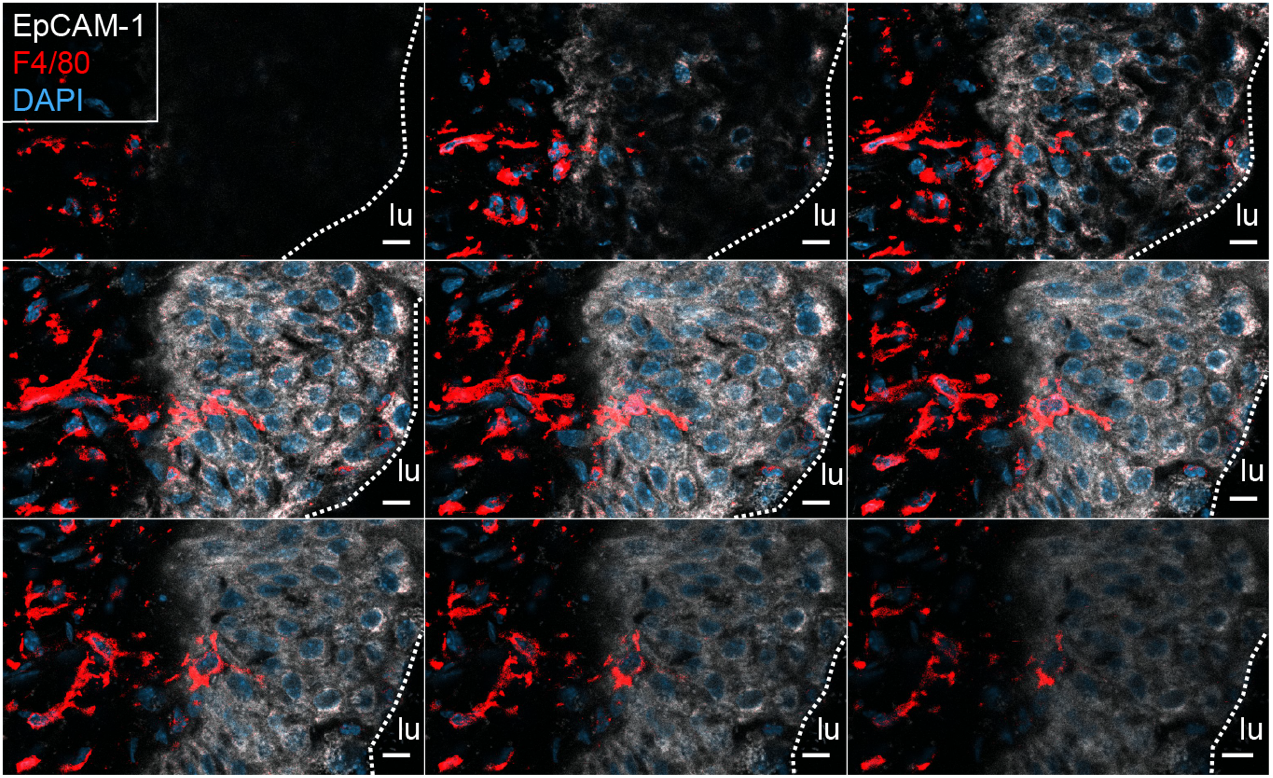
Macrophage relocation into the infected urothelium. Related to Figure 2. Bladder sections were stained with DAPI (blue), F4/80 (red) and EpCAM-1 (white) and imaged by confocal microscopy. The white dashed lines distinguish the urothelium from the connective tissue and lumen. The scale bar indicates 10 μm, step size of the Z-stacks was 0.6 μm. lu=lumen.

**Figure S3.**
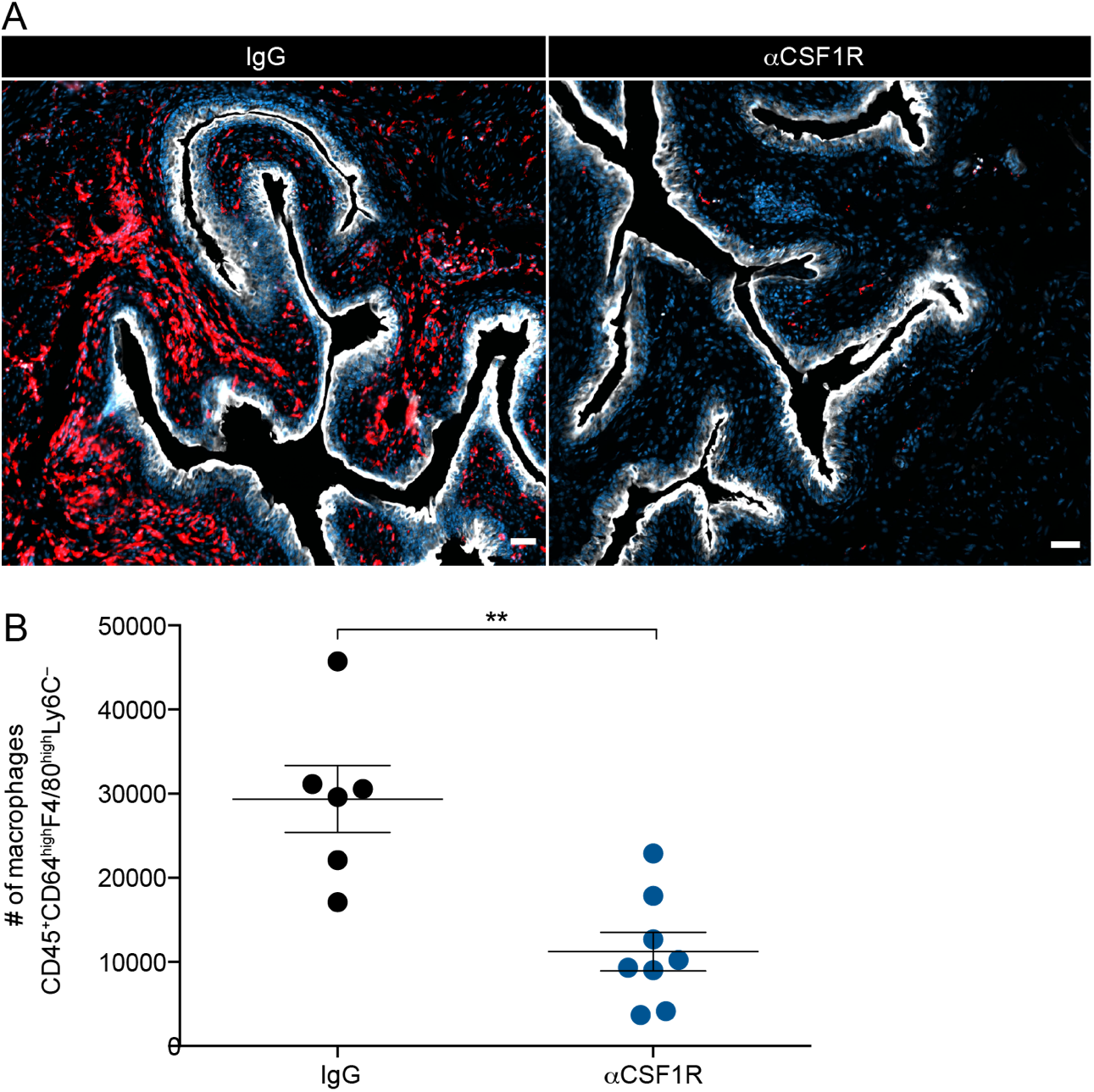
Macrophage depletion in the urinary bladder. Related to Figure 3. Female mice received two intraperitoneal injections of an aCSF1R antibody or isotype control antibodies (IgG). (**A**) Depletion of macrophages was assessed by immunofluorescence microscopy. The scale bar indicates 10 μm. (**B**) Quantification of (A). **p < 0.01. Error bars show the mean ± SEM. The scale bars indicate 50 μm.

**Figure S4.**
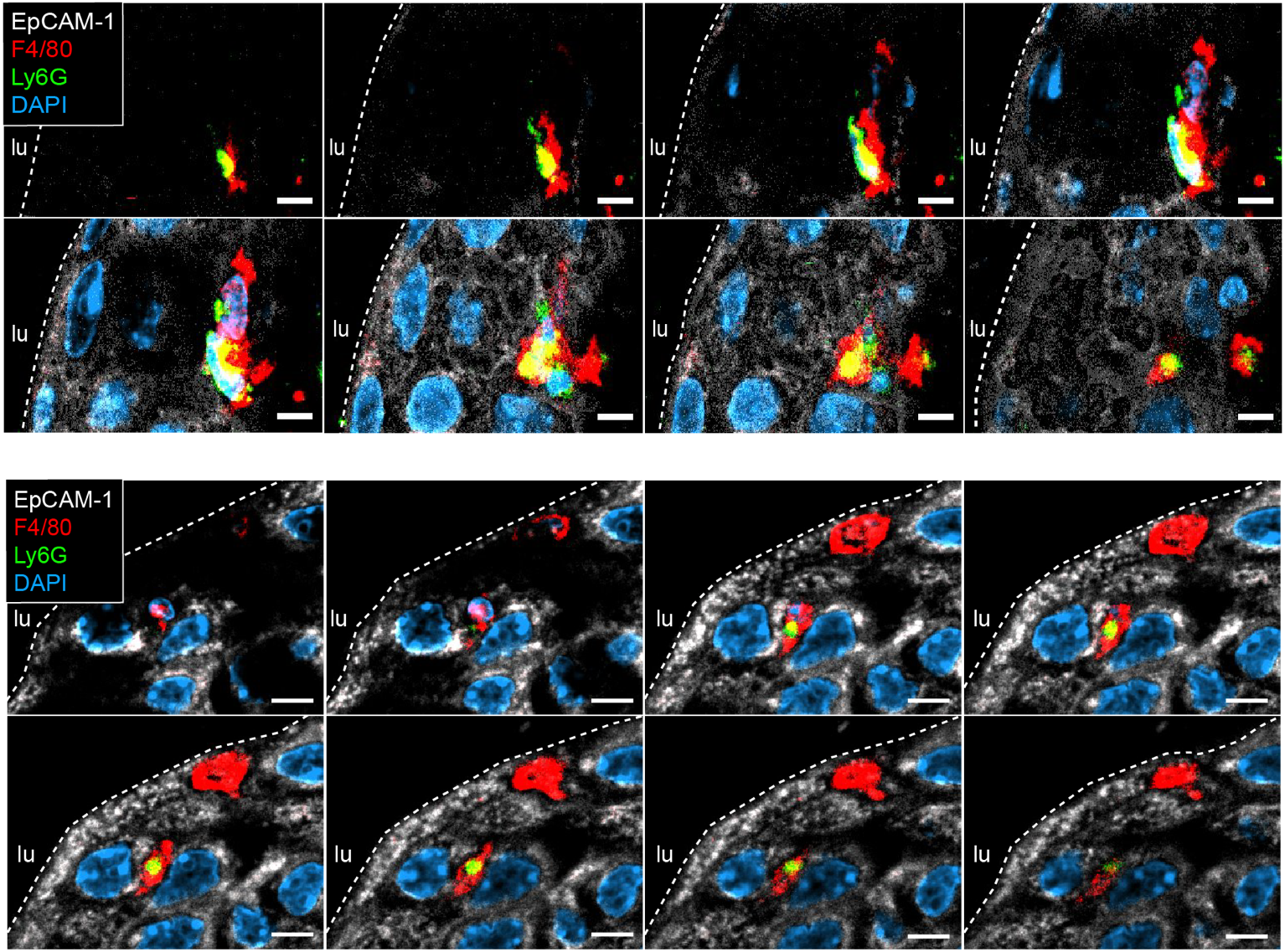
Macrophages phagocytose urothelial neutrophils upon UTI. Related to Figure 3. Bladder sections were stained with DAPI (blue), F4/80 (red), EpCAM-1 (white) and Ly6G (green) to indicate phagocytosis of neutrophils by macrophages by confocal microscopy. The scale bar indicates 5 μm, step size of the Z-stacks was 0.4 μm (top) and 0.5 μm (bottom).

